# A bistable type III secretion system is primed and loaded in *Pseudomonas aeruginosa* hypervirulent isolates with positively selected codon changes in *exsA* or *exsD*

**DOI:** 10.64898/2026.10.22.761281

**Authors:** Arianna D. Reuven, Sarah Katzenell, Alexander W. Rapp, Julian C. Oates, Joseph P. Pennington, Thomas H. Hampton, Jennifer M. Bomberger, Joshua L. Cherry, James B. Bliska

## Abstract

*Pseudomonas aeruginosa* has multiple virulence factors that promote airway infections in people with immune deficiencies. Most clinical isolates from human infections have functional type III secretion systems (T3SSs) that are subject to bistable regulation and secrete the cytotoxic effector ExoS into neutrophils. During airway infection in people with cystic fibrosis (CF), *P. aeruginosa* maintains bistable regulation or genetically adapts by inactivating or hyperactivating T3SS expression. CF clinical isolates with hyperactive T3SSs have codon changes in the transcriptional regulators, *exsA* or *exsD.* It is not known how these mutations: 1) arise, 2) promote hypervirulence, or 3) impact bistable T3SS regulation. To understand how these mutants arise, ∼24,000 genomes in the NCBI Pathogens database were searched for positively selected mutations in *exs* genes based on nonsynonomous to synonymous codon change ratios. Missense and nonsense mutations were detected in *exsA* (47) or *exsD* (45) the majority of which are predicted to be positively selected. To determine how one of these codon changes promotes hypervirulence, an *exsA*^T48I^ isolate was used to establish lung infections in mice engineered with a CF genotype. The *exsA*^T48I^ mutant and an *exsA*^+^ control had similar lung burdens, but the former caused exacerbated weight loss and lung inflammation, suggesting immunopathology from phagocyte death as the cause of hypervirulence. Two additional hyperactive mutants were identified by screening a small collection of isolates from children with CF. Immunoblotting and fluorescence microscopy showed that hyperactive *exsA* or *exsD* mutants increased production of T3SS injectisomes and effectors, indicating they have “primed and loaded” phenotypes. ExsA dimer modeling revealed that codon changes like T48I in the β-barrel domain cause a conformational change that hyperactivates the transcription factor. These results define that hyperactive mutants are positively selected, hypervirulent due to lung inflammation, and have primed and loaded T3SSs.

## Introduction

*P. aeruginosa* has multiple virulence factors and is responsible for a large percentage of airway infections that cause high morbidity and mortality in people with immune deficiencies [1]. In one key virulence mechanism, *P. aeruginosa* uses a T3SS to deliver effectors into host cells [1, 2]. The T3SS consists of an injectisome. a tip protein known as PcrV, and sets of effectors [1, 2]. Three T3SS effectors, ExoT, ExoU and ExoS are important virulence factors. All *P. aeruginosa* strains have ExoT, and ExoU or ExoS, but rarely both [3]. Population structure studies show that *P. aeruginosa* exists in two major groups, ExoS^+^ (Group A, e.g. PAO1) and ExoU^+^ (Group B, e.g. PA14) [4]. ExoS^+^ Group A strains cause most infections, including in people with cystic fibrosis (pwCF) [3, 4]. The basis for the prevalence of *exoS*^+^ *P. aeruginosa* strains in CF airway infections is unknown.

CF is an inherited disease caused by mutations in the CFTR gene which alter movement of chloride and other ions across cell membranes [5]. This alteration in ion/water flux produces highly viscous mucus and other immune deficiencies in the lungs which creates a favorable environment for opportunistic bacterial infection. CF patient mortality is often due to respiratory failure from chronic bacterial infection caused by *P. aeruginosa*.

The T3SS and ExoS are important virulence factors in acute *P. aeruginosa* lung infections [1]. In pwCF the T3SS and ExoS are likely important for establishment of new airway infections, especially in children, as these patients are often colonized with *P. aeruginosa* isolates that are T3SS positive and have antibody responses to the injectisome and ExoS [3, 6, 7]. *P. aeruginosa* isolates from established airway infections in pwCF have typically adapted genetically toward chronic persistence phenotypes such as decreased T3SS function and increased biofilm capacity [8-11]. Young pwCF can experience intermittent *P. aeruginosa* infections before age 2, and in those eligible for highly effective modulator treatment (HEMT), who experience dramatic increased health and reduced lung infection by *P. aeruginosa* as they age or in adulthood, the bacteria nevertheless persist and adapt in airways [12, 13].

ExoS has a Rho GAP domain and an arginine-specific mono ADP ribosyltransferase (ADPRT) domain. The GAP domain targets Rho GTPases, while the ADPRT domain modifies multiple proteins, such as Ras [1]. The ExoS ADPRT domain plays a major role in *P. aeruginosa* virulence by blocking phagocytosis and reactive oxygen species (ROS) production in neutrophils [14-16]. ExoS ADPRT activity is required for *P. aeruginosa* to survive in and kill neutrophils [17]. Ectopic expression studies show that ExoS ADPRT activity is directly cytotoxic to cultured mammalian cells [18]. ExoT is like ExoS in structure and required to reduce ROS and viability in neutrophils [15, 17] but less important for *P. aeruginosa* virulence in airway infections [14, 16].

Infected neutrophils can sense T3SS activity of bacterial pathogens and trigger caspase-1 inflammasome activation [19]. Insights into neutrophil caspase-1 inflammasome responses to ExoS^+^ *P. aeruginosa* T3SS have come from infection studies with laboratory strain PAO1. Flagellin secretion by the T3SS activates the NLRC4/caspase-1 inflammasome [20-23]. The NLRC4/caspase-1 inflammasome cleaves the pore-forming gasdermin D (GSDMD) protein, promoting IL-1β release and cell death (pyroptosis) from plasma membrane rupture (PMR) [21, 24, 25]. GSDMD pore formation allows Ca^2+^ influx, activating peptidyl arginine deiminase 4 (Pad4), which citrullinates histones, contributing to chromatin decondensation [21, 24]. The decondensed chromatin and citrullinated histones (e.g citrullinated histone 3, CitH3) are not extruded as neutrophil extracellular traps (NETs) [26] but instead are retained intracellularly (incomplete NET extrusion) [21, 27]. Incomplete NET extrusion is surmised to result from a restraining effect of an intact cortical actin network [21, 28]. Studies with *exoS* mutants have revealed additional layers of complexity in neutrophil responses to the T3SS. The default NLRC4/caspase-1 inflammasome response is “rewired” by ExoS ADPRT activity: ExoS inhibits NLRC4 activation and promotes GSDMD-independent PMR and CitH3 [21, 25, 27, 29]. In addition, ExoS ADPRT activity drives NLRP3/caspase-1 inflammasome activation and IL-1β release [25, 27, 29]. Controlled inflammation in response to the T3SS can be protective against *P. aeruginosa* in mouse infection models, representing a mechanism of effector triggered immunity [29-31]. Excessive inflammation can be detrimental, causing immunopathology, especially during lung infection [32], a process we refer to as effector-triggered pathogenesis [30].

T3SS function is maintained or hyperactive in some *P. aeruginosa* CF clinical isolates, and the latter phenotype can be associated with hypervirulence and lung function decline [27, 33-35]. Two *exoS^+^*CF isolates with hyperactive T3SS phenotypes resulting from codon change mutations in the negative regulator *exsD* have been identified [33, 34]. ExsD binds to and sequesters the transcription factor ExsA (**Fig.S1**) [36, 37]. ExsA is an AraC family transcription factor that acts as a dimer to drive expression of its regulon [37] (**Fig1AB.**). Codon changes in *exsD* relieve the negative regulation mechanism and result in over production and secretion of ExoS, increased *P. aeruginosa* survival in neutrophils and hypervirulence in mice [33, 34]. Jorth et al. found that a clonal isolate with a hyperactive *exsD*^T188P^ mutation was concentrated in the more damaged upper left lobe of the lung explanted from a pwCF [33]. In another study, amplicon sequencing showed that the frequency of a hyperactive *exsD*^S164P^ allele in *P. aeruginosa* clonal isolates increased during a three-year period of lung decline in a pwCF [34]. Subsequently, a T3SS-inactivating mutation (*exsA*^Q83*)^ was detected in *P. aeruginosa* isolates from this pwCF [34].

We identified an *exoS^+^ P. aeruginosa* strain with a hyperactive T33S that appeared transiently among a collection of isolates collected longitudinally from the sinuses of a young adult pwCF [27, 35]. This isolate, p32_85, has a codon change in *exsA* (T48I, polar to hydrophobic) [27, 35]. T48I is a solvent exposed residue on a strand of the β-barrel domain, a part of the N-terminal regulatory region of ExsA that interacts with ExsD [36, 38-40]. However, ExsA^T48I^ shows no defect in ExsD interaction as shown by bacterial two-hybrid analysis [27]. The *exsA*^T48I^ mutant promotes enhanced CitH3 and PMR in infected neutrophils [27]. The ninjurin-1 protein (NINJ1) [41-44] was implicated in ExoS-driven PMR in neutrophils infected with the *exsA*^T48I^ mutant [27]. These data support the idea that codon changes that activate or inactivate the T3SS can arise in *P. aeruginosa exsD* or *exsA* in a location and time dependent manner and impact disease during airway infection in CF. However, the prevalence of hyperactive strains is unknown and the mechanisms of their hyperactivation and hypervirulence remain to be fully defined.

Isogenic *P. aeruginosa* populations exposed to an activating signal (e.g. low Ca^2+^) exhibit phenotypic heterogeneity with respect to T3SS expression [37, 45-47]. A fraction of *P. aeruginosa* cells in the population highly express T3SS genes (on) and the remainder do not (off) (**Fig.S1**), a phenomenon known as bistability [48]. Bistable expression of the T3SS allows *P. aeruginosa* to balance the benefits and costs of this virulence apparatus [45, 48, 49]. Bistability of the *P. aeruginosa* T3SS is controlled by ExsA, ExsD and two other products of the *exsCEBA* operon, ExsC and ExsE (**Fig.S1, Fig.1AB**). ExsD, ExsE and ExsC work in a partner switching mechanism to control ExsA activity (**Fig.S1**). In response to activating signals, ExsE is secreted through the injectisome, releasing ExsC to bind ExsD and free up ExsA to drive maximal T3SS gene expression (on)(**Fig.S1**). Recent work suggests that in the off conformation ExsA expression is below the limit of detection even in the presence of activating signal [45, 49]. In addition, expression of ExsA from its own promoter under control of cyclic AMP (cAMP) and the transcription factor Vfr (**Fig.1A**) results in free ExsA that can prime *P. aeruginosa* to express and assemble injectisomes in the absence of activating signal (primed, **Fig.S1**)[45, 47, 49]. Notably, *P. aeruginosa* Δ*exsD* or *exsD*^T188P^ mutants express effectors in the absence of activating signal [33, 50]. A key observation is that nearly all the bacteria in a *P. aeruginosa* Δ*exsD* mutant population are primed for T3SS expression [45].

**Fig.1.**
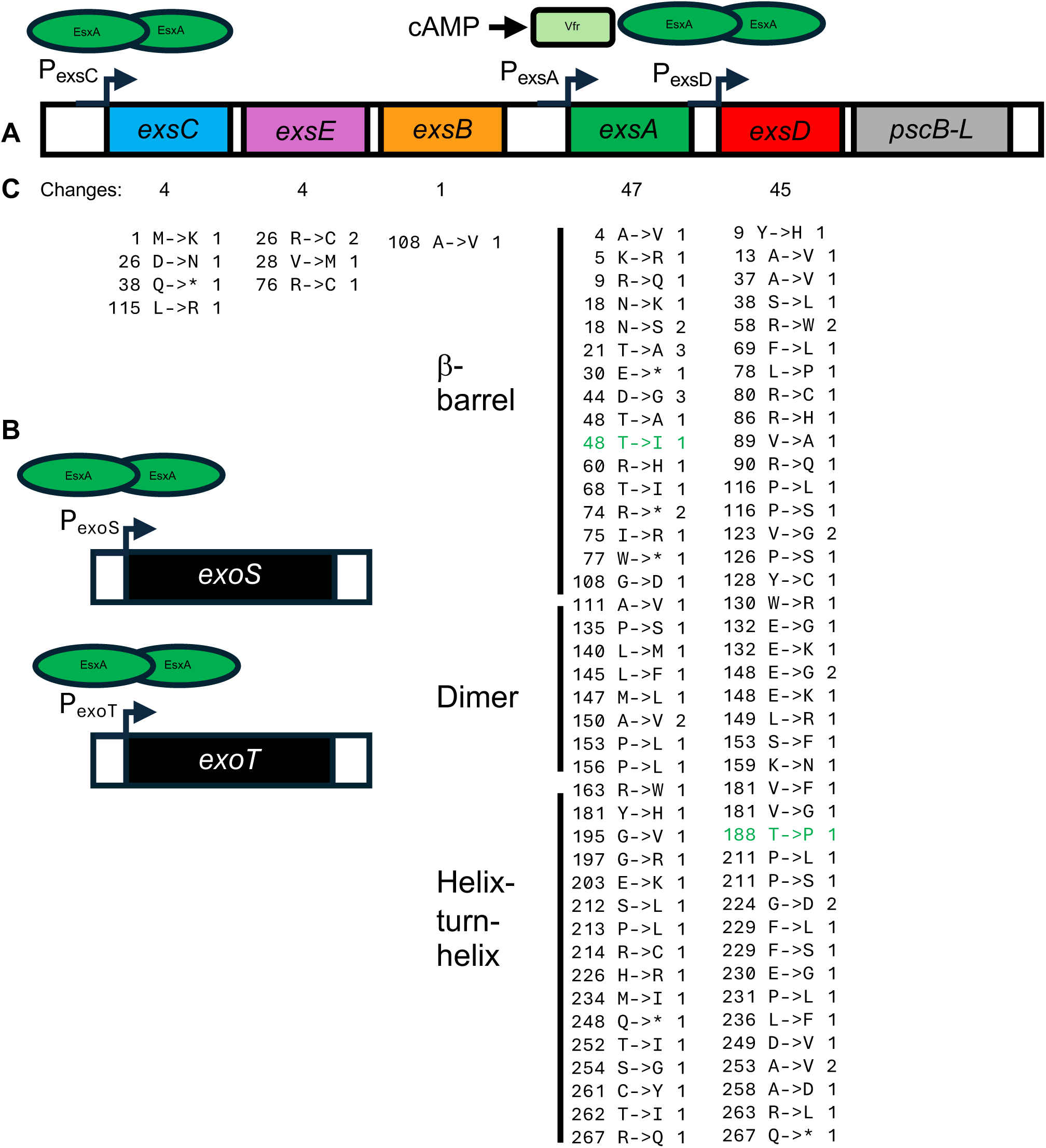
P. *aeruginosa* ExsA regulon and codon changes in *exs* genes identified in sequenced isolates. ExsA as a dimer activates transcription of (A) the *exsCEBA* operon and the *exsDpscB-L* operon (and other T3SS operons not shown) and (B) the *exoS* and *exoT* genes. ExsC, ExsE and ExsD regulate ExsA activity, PscB-L contribute to assembly of the T3SS injectisome, and ExoS and ExoT are effectors. Vfr activated by cAMP drives transcription of *exsA* from a second promoter. (C) The *exs* genes in ∼24,000 sequenced isolates in the NCBI Pathogens database were subjected to a stringent search for codon changes to identify potential sites of positive selection. The numbers of missense or nonsense (*) codon changes identified are shown below each gene. Lists show changes by amino acid position, change, and number of independent occurrences. Isolates with codon changes in green been previously identified, published and demonstrated to have hyperactivating mutations, respectively. Lines to the left of the *exsA* list demark the positions of codon changes with respect to the three different functional domains of ExsA.

Here we investigated the origin and prevalence of *P. aeruginosa* hyperactive T3SS mutants in genome and strain collections and defined how positively selected codon changes in the ExsA β-barrel domain impact dimer conformation and hypervirulence in a CF mouse lung infection model. In addition, we investigated how hyperactivating codon changes in *exsA* or *exsD* impact T3SS bistability regulation.

## Results

### Identification of positively selected codon changes in *exs* genes in sequenced *P. aeruginosa* isolates

Gain or loss of function codon changes in *exsA* or *exsD* of *P. aeruginosa* can arise during airway infection in CF and can be associated with disease [27, 33, 34]. Questions remain about how prevalent these mutations are and if they are positively selected. To address these questions, we analyzed clusters of closely related *P. aeruginosa* genomes in the NCBI Pathogens database to identify recent changes in the five *exs* coding sequences (**Fig.1A**). An excess of missense or nonsense changes relative to a neutral expectation can, much like a dN/dS value greater than one, provide evidence for positive selection. Previous analysis of *Salmonella* genomes identified hundreds of genes with signatures of positive selection that result in gain or loss of function codon changes [51, 52]. Of note, activating mutations are enriched in genes involved in transcriptional regulation in *Salmonella* [52]. Inactivation of one of the genes identified, *glpT*, was shown to confer an advantage during the early stage of *Salmonella* Typhimurium infection but a disadvantage during later stages, which may be regarded as a case of antagonistic pleiotropy [53].

Changes in the *P. aeruginosa exs* genes were inferred from a total of ∼24,000 mostly clinical isolates. To increase the stringency of the search, we focused on results that met strict requirements for aligned raw sequence reads and were derived from a cluster of at least 5 isolates. The results of this analysis are shown in **Fig.1C**.

For *exsA* and *exsD*, the tallies suggest positive selection for nonsynonymous changes. For *exsA*, the ratio of missense to synonymous changes, 42:5=8.4, exceeds the estimated ratio of rates under selective neutrality (2.6) by a factor of 3.2. It can also be estimated directly that only 7.1 missense changes are expected under selective neutrality. Thus, the majority of the observed missense changes are likely the result of positive selection. For *exsD*, the missense:synonymous ratio, 44:3=14.7, also greatly exceeds the neutral prediction (2.5), as does the absolute number of missense changes (44 observed vs. 6.4 expected), pointing to a similar conclusion. The five changes producing premature stop codons in *exsA* also significantly exceed the neutral expectation, suggesting positive selection for inactivation of *exsA* under some conditions.

Notably, the published *P. aeruginosa* CF clinical isolate with the *exsA*^T48I^ hyperactivating [27, 35] mutation was identified in this analysis (**Fig.1C**). A second isolate with a change in *exsA* at position 48 (T48A) was identified (**Fig.1C**). Several positions were changed in multiple independent clinical airway isolates (e.g. N18, **Fig.1C**, **Table 1**). In addition, the published *P. aeruginosa* CF clinical isolate with a hyperactivating T188P mutation [33] was also represented among the 45 nonsense and missense codon changes in *exsD* (**Fig.1C**). Many of the isolates with codon changes in *exsA* or *exsD* were from airway infections including CF (**Table 1**), but also from a range of other human sites including blood stream and urinary tract. These results suggest that *exsA* and *exsD* are hot spots for positively selected hyperactivating or loss of function codon changes.

**Table 1.** Examples of *exsA* β-barrel codon changes in *P. aeruginosa* airway isolates.

|  |  |  |  |  |
| --- | --- | --- | --- | --- |
| Codon change | Source | #Clonal isolates | Hyperactive (expected) | Biosample |
| N18S (AAC->AGC) | CF infant bronchiectasis | 13 | (Yes) | SAMN14160409 |
| T48I (ACT->ATT) | CF | 2 | Yes | SAMN20695873 |
| T68I (ACC->ATC) | Bronchiectasis | 16 | (No) | SAMEA114323275 |

For *exsE*, four missense and no synonymous changes were observed (**Fig.1C**). These numbers do not allow any definite conclusions, but the nature of the amino acid changes is telling. Binding of ExsE to ExsC involves two Arg-X-Val-X-Arg motifs [54]. All four nonsynonymous changes in *exsE* are in these motifs and eliminate either the first arginine, which forms multiple salt bridges with ExsC, or the valine. These changes are expected to disrupt binding of ExsE to ExsC, leading to a hypersecretory T3SS phenotype, and presumably were selected for that reason. Of the three missense and one nonsense changes detected in *exsC* (**Fig.1C**), the M1K and Q38* mutations would be expected to decrease or eliminate full length protein production, resulting in loss of T3SS function. A single conservative A108V change was observed in *exsB* (**Fig.1C**).

### The *P. aeruginosa exsA*^T48I^ mutant is hypervirulent in CF mouse lung infection model

To determine if the isolate with the T3SS hyperactivating codon change *exsA*^T48I^ (p32_85, **Table 2**) [27, 35] is hypervirulent, we infected C57BL/6 mice engineered to have a CF genotype (F508del) [55]. We used a sublethal infection dose to follow weights and lung CFU up to 5 days post infection. Lung infections in F508del mice were established with the isogenic control p32_08 (*exsA*^+^) (**Table 2**) or p32_85 (*exsA*^T481^), followed by survival, weight, and lung CFU and H&E assays. All mice survived but those infected with the hyperactive isolate had significantly lower weights than the controls between 36 and 120 h (**Fig.2A**). At 120 h the lung CFUs were not significantly different (**Fig.2B**). Representative lung H&E images suggested increased inflammatory cell recruitment in response to the hyperactive isolate as compared to p32_08 (*exsA*^+^) (**Fig.2C, Fig.S2**). These data indicate that the *exsA*^T481^ isolate is hypervirulent, causing more inflammation and disease at similar organ burdens as the control.

**Fig.2.**
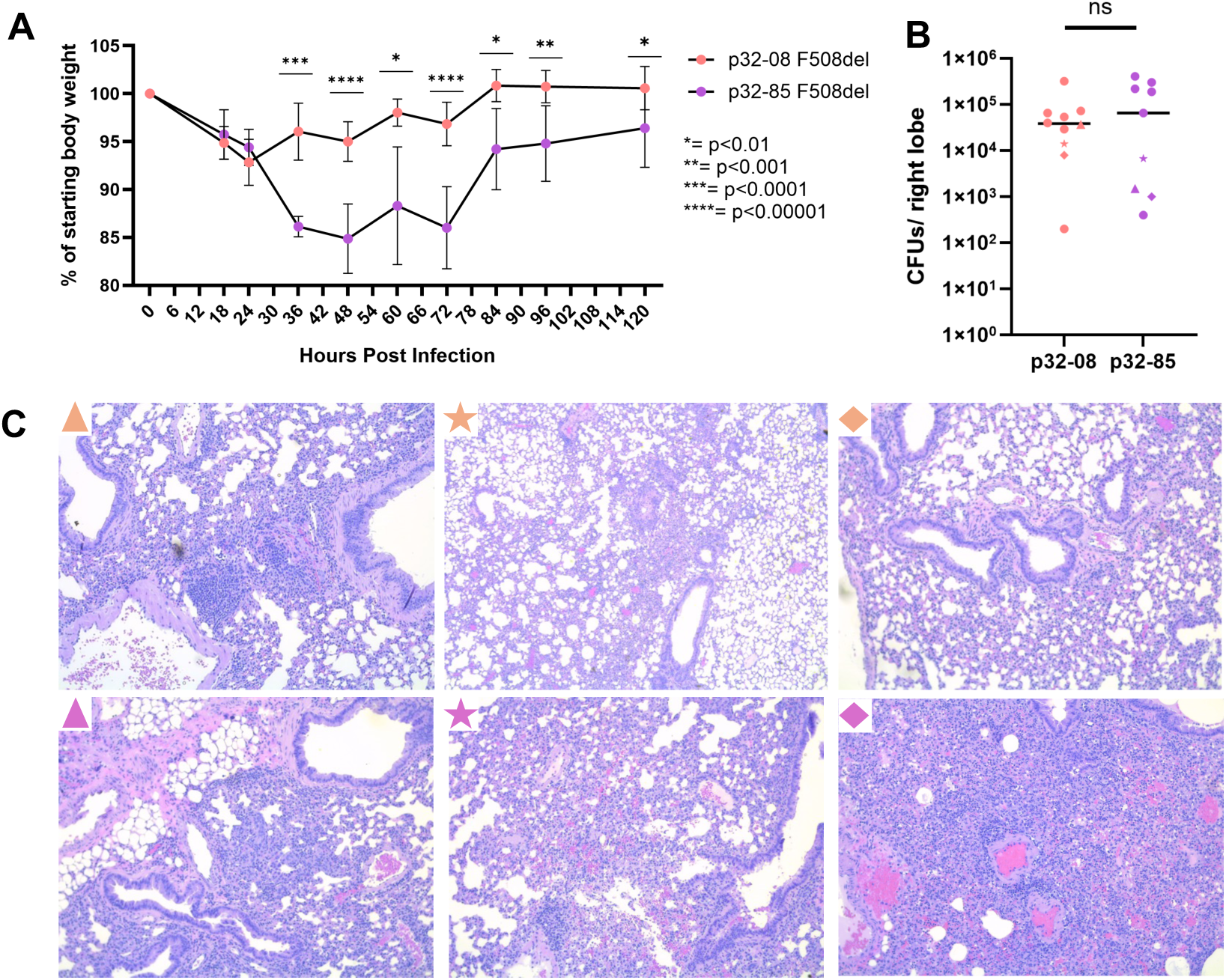
Outcomes of F508del mouse infection with p32 isolates 08 (*exsA*^+^) or 85 (*exsA*^T48I^). Groups of F508del female and male mice were infected OP with 5x10^5^ CFU of p32_08 or p32_85. Weight and CFU data was pooled from 3 independent experiments, n=10 and 9, respectively. (A) Percent of starting body weight over time. Error bars are SD. Differences between weights that were significant by multiple T-test and a multiple-comparisons correction (Holm–Šídák) are shown. (B) Right lung lobe CFU at 120 h, bars are average, p=not significant by Mann Whitney test. Triangle, star and diamond symbols correspond to lungs that were analyzed by H&E. (C) Representative H&E images of left lobe lung sections at 120 h and 10x magnification.

**Table 2.** *P. aeruginosa* strains used in this study.

| Table 2. *P. aeruginosa* strains used in this study |  |  |  |
| --- | --- | --- | --- |
| Name | Relevant genotype | Description | Ref |
| p32_08 | *exsA*<sup>+</sup> | CF isolate | [35] |
| p32_08 Δ*ExsA* | Δ*exsA* | Deletion mutant of p32_08 | This study |
| p32_08 *ExsA*<sup>T48I</sup> | *exsA*<sup>T48I</sup> | Allele swap mutant of p32_08 | [27] |
| p32_85 | *exsA*<sup>T48I</sup> | CF isolate | [35] |
| p32_85 *ExsA*<sup>+</sup> | *exsA*<sup>+</sup> | Allele swap mutant of p32_85 | [27] |
| PAO1F |  | Laboratory strain | [17] |
| PAO1F Δ*pscD* | Δ*pscD* | Deletion mutant of PAO1F | [17] |
| DH1137 |  | CF isolate (originally called NC-AMT0101-2) | [70] |
| 31-1 |  | CF isolate | [59] |
| 33-1 | *exsA*<sup>E104G</sup> *exsD*<sup>Q241*</sup> | CF isolate | [59] |
| 36-2 |  | CF isolate | [59] |
| 41-1 |  | CF isolate | [59] |
| 47-2 |  | CF isolate | [59] |
| 60-2 |  | CF isolate | [59] |
| 62-1 | *exsD*<sup>R80S S105A</sup> | CF isolate | [59] |
| 66-1 |  | CF isolate | [59] |
| 71-1 |  | CF isolate | [59] |
| 74-1 |  | CF isolate | [59] |
| LU *ExsD*<sup>T188P</sup> | *exsD*<sup>T188P</sup> | CF isolate | [33] |

### ExsA^T48I^ promotes a “primed and loaded” T3SS phenotype in *P. aeruginosa*

*P. aeruginosa* hyperactive *exsD* mutants display increased production and secretion of ExoS under T3SS non-activating or activating conditions, respectively [33, 34, 50]. In addition, a *P. aeruginosa* Δ*exsD* mutant appears to be fully primed in the absence of activating signal as shown using a fluorescent reporter under control of the *exoT* promoter [45]. We used immunoblotting to determine if ExsA^T48I^ increases production of T3SS gene products under non-activating or activating conditions. *P. aeruginosa* p32_08 (*exsA*^+^) or p32_85 (*exsA*^T48I^) were grown in the presence of EGTA (low Ca^2+^ activating conditions) and lysates immunoblotted for ExsA. Production of ExsA^T48I^ was ∼2 fold reduced as compared to ExsA (**Fig.3A**, lane 3 vs 2). ExsA production was restored to normal levels in an p32_85 mutant in which *exsA*^T48I^ was reverted to *exsA*^+^ (lane 4). The p32 strains were next grown in the presence or absence of EGTA and lysates immunoblotted for ExsD, ExoS or PcrV. ExsD was produced at lower levels in both strains under the -EGTA conditions (**Fig.3B**, lanes 4,5 vs 2,3). In addition, under the -EGTA conditions there was ∼7 fold more ExsD produced by ExsA^T48I^ vs ExsA^+^ (lane 5 vs 4). ExoS was produced at lower levels in the *exsA*^+^ background under the -EGTA conditions (**Fig.3C** top panel, lane 2 vs 5). In contrast, ExoS was produced at similar levels in the *exsA*^T48I^ background under both conditions (lane 3 vs 6). The production of PcrV followed a similar pattern as ExoS (**Fig.3C**, third panel). No ExoS or PcrV was produced by a p32_08 Δ*exsA* mutant (**Table 2**) analyzed in parallel as a negative control (**Fig.3C**, lanes 4,7). Analysis of the secreted fraction showed that ExoS was not exported by the hyperactive *exsA*^T48I^ mutant under the -EGTA conditions (**Fig.3C** second panel, lane 6). In contrast, PcrV was exported by the hyperactive *exsA*^T48I^ mutant under the -EGTA conditions (**Fig.3C** fourth panel, lane 6).

**Fig.3.**
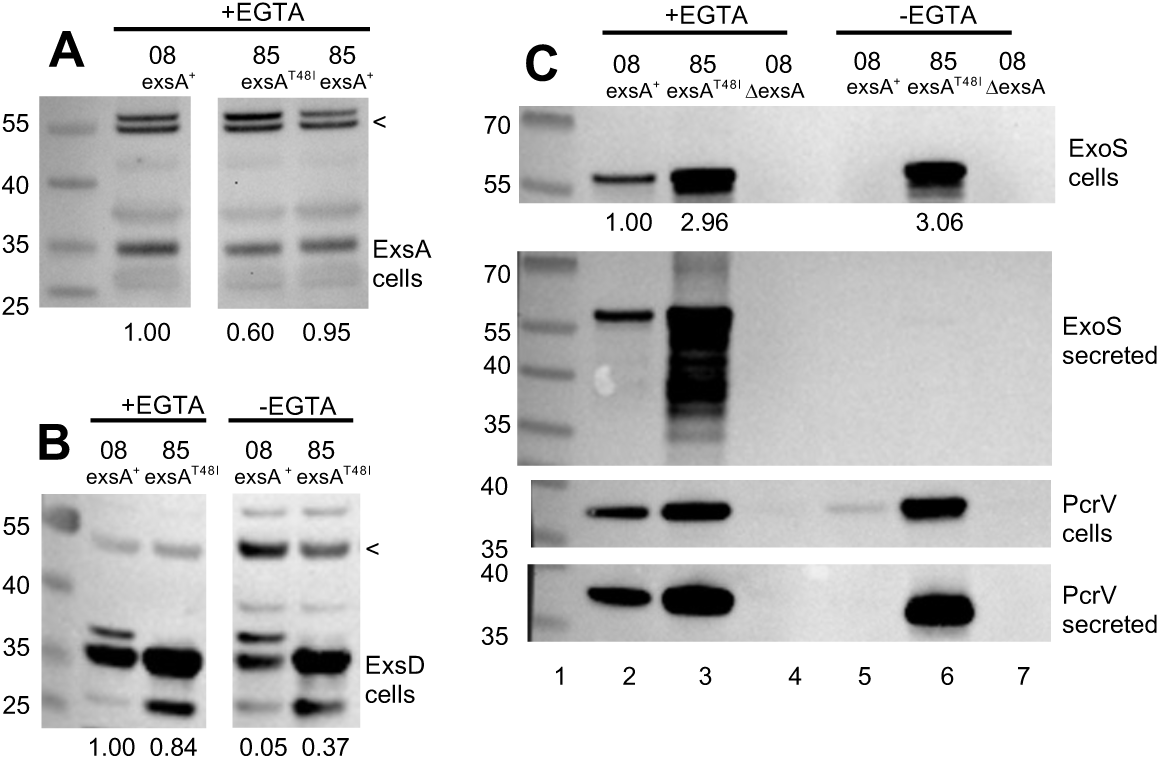
Immunoblot analysis of ExsA, ExsD, ExoS and PcrV production or secretion in p32 strains. Bacterial cell lysates (A, B, C first and third) or secreted fraction (C second and fourth) of the indicated P32 strains grown in LB under low Ca^2+^ (T3SS activating) or high Ca^2+^ (T3SS non-activating) conditions for ∼3 h were immunoblotted for (A) ExsA, (B) ExsD or (C) ExoS. The first lane in each panel is MW standards. Non-specific bands (<) in A,B were used as loading controls for quantifying ExsA or ExsD bands, normalized to low Ca^2+^ 08 *exsA*^+^, as shown at the bottom of each lane. In (C) ExoS lysate bands were quantified and normalized to low Ca^2+^ 08 *exsA*^+^, as shown at the bottom of each lane.

Fluorescence microscopy was used to compare bistability phenotypes in p32_08 and p32_85 at the single cell level. A super folding (sf) GFP transcriptional reporter under control of the *exoT* promoter [45] was introduced into the p32_08 (Δ*exsA* or *exsA^+^*) or p32_85 (*exsA*^T48I^) isolates. The bacteria were grown in -/+EGTA conditions, followed by fixation and staining for total bacteria (Hoechts), and images were captured by fluorescence microscopy and quantified. **Fig.4A** shows representative images and **Fig.4B** has quantification of mean fluorescence intensity (MFI). In the absence of EGTA p32_08 Δ*exsA* and *exsA^+^* bacteria had little to no GFP signal and similar MFI, reflecting background, which was confirmed by imaging p32_08 lacking the reporter (**Fig.4AB**). Compared to p32_08 Δ*exsA,* p32_08 *exsA^+^* grown in the presence of EGTA displayed GFP^+^ signal in a fraction of the bacteria reflecting the bistable T3SS “on” (T3SS^+^) phenotype (**Fig.1**, **Fig.4A**). Using p32_08 Δ*exsA* for background correction, we measured the on fraction for each strain in -/+EGTA with a false-positive rate of 0.01%. The on-fraction for p23_08 went from 0% to 10.21% (**Fig.4AB**). In the absence of EGTA, a greater fraction of p32_85 bacteria were GFP^+^ and had significantly higher MFI than p32_08 *exsA^+^* (**Fig.4AB**), confirming the ExoS immunoblot results (**Fig.3C**) indicating that the *exsA*^T48I^ mutation causes a “primed and loaded” T3SS phenotype. Activation of the T3SS with EGTA significantly increased the MFI in p32_85 compared to -EGTA and the on fraction went from 47% to 89% (**Fig.4AB**) indicating that the *exsA*^T48I^ mutation does not result in a fully primed phenotype.

**Fig.4.**
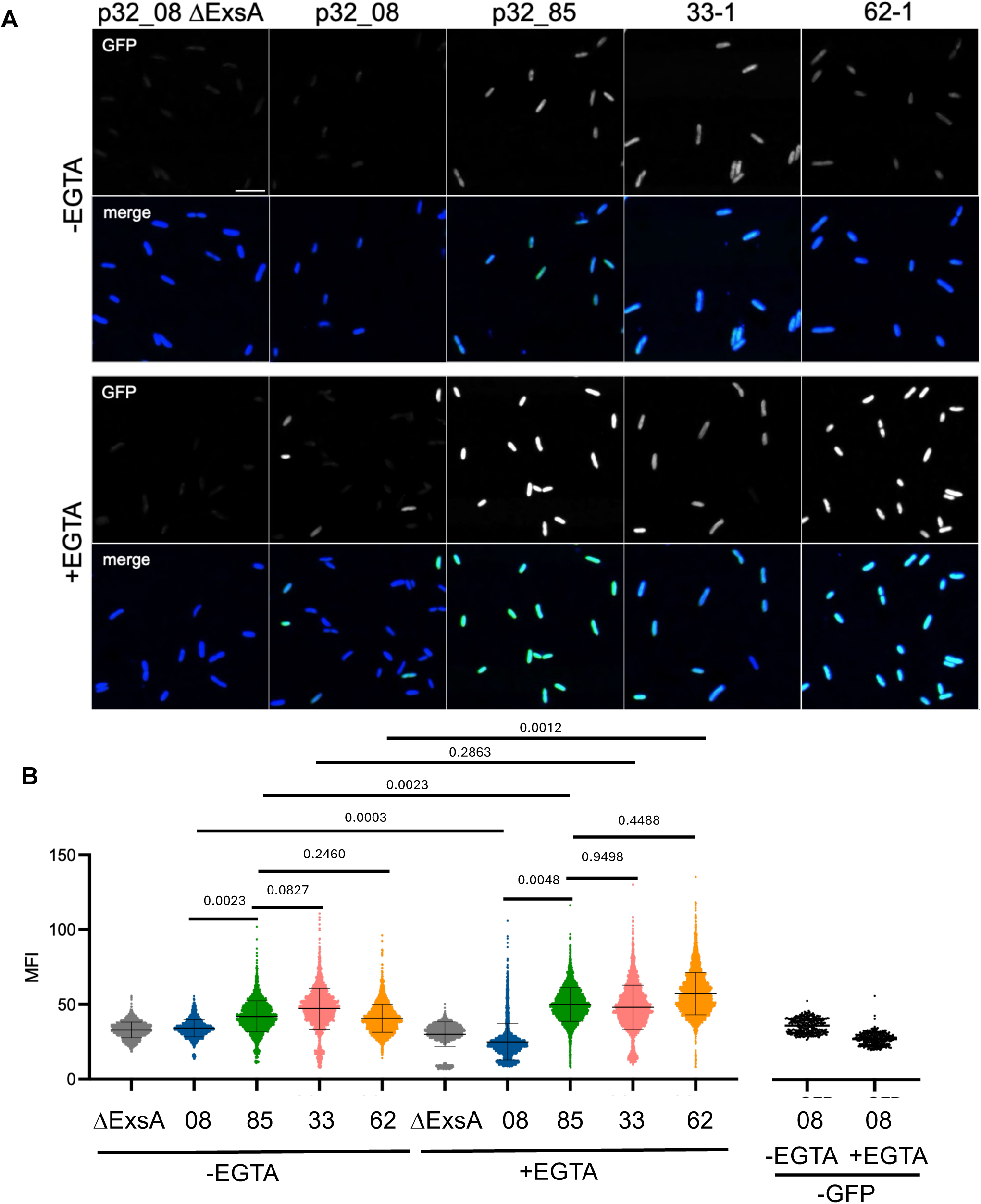
Microscopic analysis of T3SS bistability phenotypes in *P. aeruginosa* using pexoT-GFP reporter. The indicated strains with the p*exoT-*GFP reporter were grown in in absence or presence of EGTA media to induce T3SS expression, followed by fixation and Hoechst staining, and images were captured by fluorescence microscopy. (A) Representative images from a single experiment of GFP^+^ bacteria and merged GFP^+^ (green) and total (blue) bacteria. Bar is 5 microns. (B) The mean fluorescence intensity (MFI) of individual bacteria pooled from 4-5 experiments. Groups were compared by a mixed model paired t-test with Bonferroni-Holm correction for multiple comparisons. Control data from p32_08 with no GFP reporter is from a single experiment.

In summary, these results suggest that as compared to ExsA^+^, ExsA^T48I^ does not substantially increase its own production, but does increase production of ExsD, ExoS and PcrV, especially under -EGTA non-activating conditions. Under -EGTA non-activating conditions PcrV is secreted, indicating that ExsA^T48I^ increased priming (production of injectisomes on the bacterial surface). In addition, ExsA^T48I^ increased production but not secretion of ExoS in the absence of EGTA, indicating that the hyperactive isolate is “primed and loaded”.

Expression of the flagellar T3SS is inversely regulated with the T3SS injectisome [37, 56]. To determine if the hyperactive *exsA*^T48I^ mutant p32_85 has reduced flagellar motility, swimming assays were performed. The p32_08 or p32_85 strains were grown in the absence or presence of EGTA and spotted onto swim agar plates. As shown in **Fig.S3A**, irrespective of EGTA, p32_08 had substantially increased motility as compared to p32_85, consistent with the primed and loaded hyperactive T3SS in the later isolate. To confirm that the motility phenotype was specific to the *exsA*^T48I^ mutation, we assayed p32_08 and p32_85 “allele swap” strains (**Table 2**). As shown in **Fig.S3B**, the *exsA*^T48I^ mutation caused a significant reduction in motility in both strain backgrounds.

### Evidence that T48I induces a conformational change in ExsA

A large number of missense codon changes identified in *exsA* correspond to the β-barrel (18 total shown in **Fig.2C**, 12 positions shown in **Fig.5A**). The codon changes are not restricted to one region of the β-barrel (**Fig.5A**), suggesting the entire domain may be sensitive to amino acid substitutions that activate or inactivate ExsA. The β-barrel of the AraC transcription factor regulates a dimer conformational change in response to binding of arabinose [57]. However, the role of the β-barrel in regulating ExsA is unknown. Activation of ExsA-dependent promoters requires the successive binding and dimerization of ExsA at two adjacent sites (sites 1 and 2) near the −35 region [37]. To investigate if T48I causes a conformational change in ExsA a dimer model was created using AlphaFold 3 (AF3) (**Fig.5B**) [58]. Like a dimer model based on the crystal structure of the ExsA N-terminal region [39] the helix-turn-helix (HTH) domains that bind DNA are on opposite sides of the WT dimer and positioned close to each β-barrel (**Fig.5B**). In contrast, the ExsA^T48I^ dimer shows a dramatic conformational change in which flexible linker regions allow the two HTH domains to swing down into closer proximity to each other (**Fig.5C**). In AF3 the Predicted Template Modeling Score (pTM) measures the accuracy of the overall predicted structure of an individual chain or a whole complex. Scores above 0.5 indicate a likely correct overall fold. The pTM values for dimer structures of ExsA and ExsA^T48I^ are 0.66 and 0.55, respectively. It is unclear if the HTH domains in the AF3 model of the ExsA^T48I^ dimer are ideally positioned to bind sites 1 and 2 of canonical promoters, as answering this question will require additional modeling with DNA. This result does raise the possibility that the β-barrel, by an unknown mechanism, possibly by binding an unidentified small molecule, regulates a conformational change in ExsA dimers that allows binding to sites 1 and 2 in promoters (**Fig.5D**). We hypothesize that ExsA^T48I^ is constitutively active for this conformational change, resulting in the hyperactive T3SS phenotype (**Fig.5E**). In a preliminary test, we prioritized two *exsA* codon changes, N18S and T68I, for dimer conformation analysis by AF3 (**Table 1**). N18S and T68I occurred in airway infections and were found in many clonal isolates (**Table 1**). Interestingly, N18S showed the ExsA dimer conformation change, while T68I did not (**Fig.S4**). The pTM value for the dimer structure of ExsA^N18S^ was 0.57.

**Fig.5.**
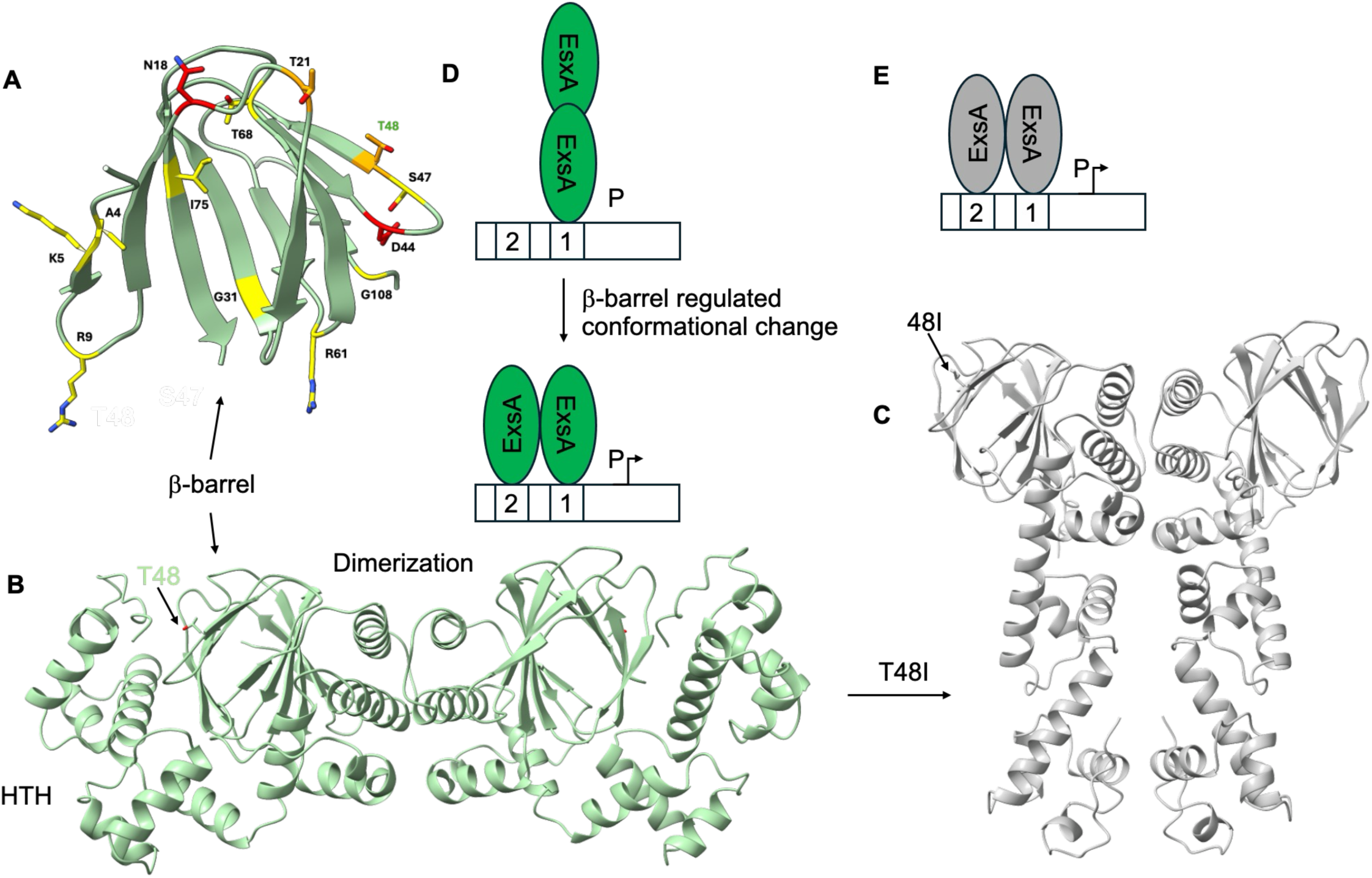
AlphaFold 3 (AF3) models of ExsA, ExsA^T48I^ and dimer conformation changes. (A) AF3 of ExsA β-barrel residues 1-80 showing 12 positions identified as potential sites of positive selection as shown in Fig.1C, highlighted in yellow for one, orange for two or red for three independent occurrences. (B) AF3 model of ExsA dimer in green showing locations of β-barrel and T48, dimerization and helix-turn-helix (HTH) regions for the left monomer. (C) AF3 model of ExsA^T48I^ dimer in grey showing conformational change in locations of HTH domains. (D) Model of ExsA head-to-head dimer interacting via one HTH with promoter (P) initially at Site1. Following a conformational change regulated by the β-barrel, Site2 can be occupied by the second HTH, leading to promoter activation. (E) Model of ExsA^T48I^ dimer that constitutively adopts a conformation that can occupy both sites and drive transcription. In AF3 the Predicted Template Modeling Score (pTM) measures the accuracy of the overall predicted structure of an individual chain or a whole complex. Scores above 0.5 indicate a likely correct overall fold. The pTM values for dimer structures of ExsA and ExsAT48I are 0.66 and 0.55, respectively.

### Identification of *P. aeruginosa* hyperactive T3SS isolates from children with CF

To test if hyperactive T3SS isolates could be identified by screening banked CF clinical strains, we analyzed a collection of 10 strains from children with CF of less than 11 years of age (**Table 2**) [59]. The strains were grown in +EGTA conditions and secretion of ExoS and PcrV were detected by immunoblotting. For controls we used PAO1F, a *pscD* mutant of PAO1F, and a clinical isolate (DH1137) that has low T3SS activity [60] (**Table 2**). As shown in **Fig. 6A**, 5 out of 10 isolates secreted ExoS or PcrV at near to or above the level of the positive control PAO1F (33-1, 44-1, 62-1, 71-1 and 74-1). The isolate 71-1 was unusual in that PcrV but not ExoS was secreted (lane 12). Since 33-1 and 62-1 appeared to be T3SS hyperactive, the *exsA* and *exsD* genes from these isolates were sequenced. As compared to PAO1, 33-1 has E104G and Q241* codon changes in *exsA* and *exsD*, respectively (**Table 2**). The E104G change is non-conservative and located in the β-barrel domain of ExsA and could play a role in hyperactivation. The Q241* change is predicted to truncate ExsD and likely causes T3SS hyperactivation in this strain. Strain 62-1 has R80S and S105A codon changes in *exsD* (**Table 2**), either of which could be responsible for hyperactivation. Immunoblotting of bacterial cell lysates showed that 33-1 produces a truncated ExsD protein as expected from the Q241* mutation (**Fig.6B**). To confirm that 33-1 and 62-1 have hyperactive “primed and loaded” T3SS phenotypes, the strains were grown in -EGTA or +EGTA conditions and secreted and cell lysate samples were immunoblotted for ExoS and PcrV. Both strains produced ExoS and PcrV irrespective of EGTA and only hypersecreted ExoS in +EGTA (**Fig.6C**). Notably, both strains secreted PcrV irrespective of EGTA (**Fig.6C**). Fluorescence microscopy analysis of 33-1 and 62-1 containing the p*exoT*-sfGFP reporter grown in the absence of EGTA confirmed a hyperactive “primed and loaded” T3SS phenotype similar to p32_85 (**Fig.4AB**). The MFI of 33-1 did not significantly increase between -EGTA and +EGTA (**Fig.4B**) with the on fraction changing from 82% to 86%, suggesting this strain has a fully primed T3SS phenotype, likely as a result of the loss of function *exsD*^Q241*^ mutation. Like p32_85 the MFI of 62-1 significantly increased when shifted from -EGTA to +EGTA (**Fig.4B**), with the on fraction going from 28% to 97%, indicating this strain does not have a fully primed phenotype.

**Fig.6.**
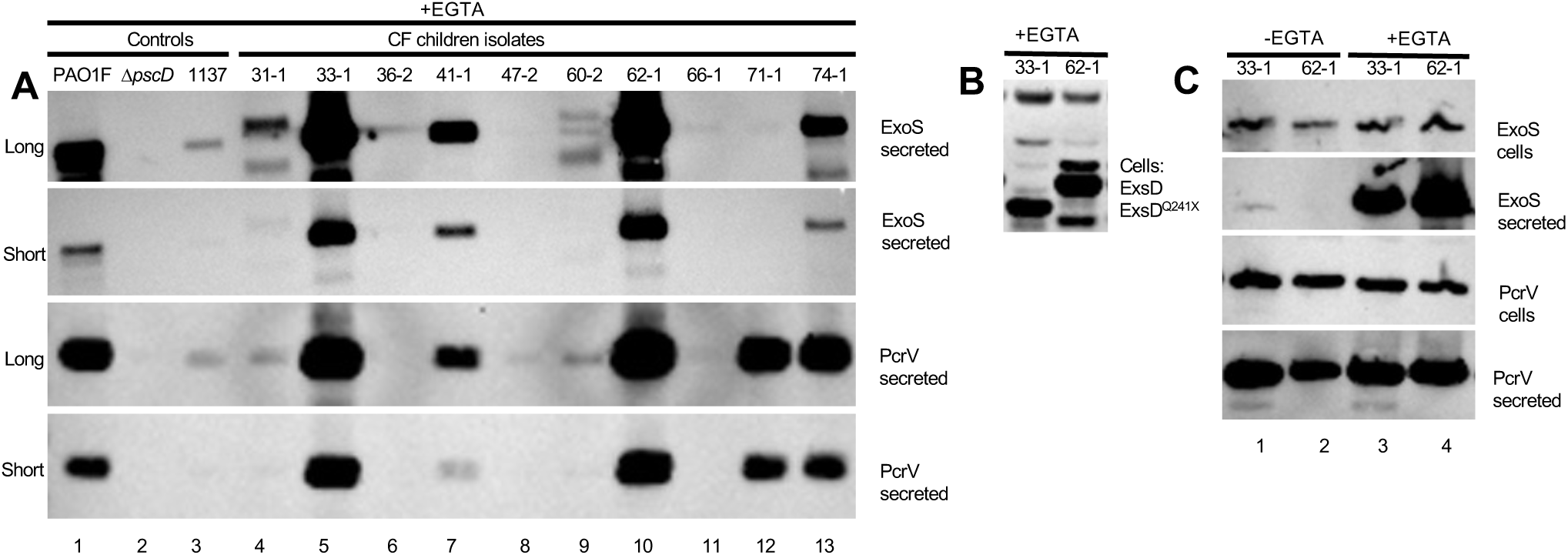
Analysis of ExoS, PcrV and ExsD production or secretion in *P. aeruginosa*. Bacterial cell lysates or secreted fractions of the indicated *P. aeruginosa* strains grown in presence or absence of EGTA for 3 h were immunoblotted with antibody to (A, D) ExoS and PcrV, or (C) ExsD. In (A) laboratory strain PAO1F, a T3SS negative mutant (Δ*pscD*) and mucoid isolate DH1137 served as controls in comparison to the 10 CF children isolates. Long and short exposures of the immunoblots are shown. In (B) the codon changes in *exsA* or *exsD* detected in 33-1 or 62-1 by sequencing are shown.

To determine if 33-1 and 62-1 have enhanced PMR activity, we carried out BMN infections. BMNs were purified from C57BL/6J mice and stimulated with low dose LPS, which extends their lifespan [61, 62] and primes inflammasome responses [27]. Flow-cytometry on primed BMNs confirmed their purity and viability (∼95%) [27]. BMNs were infected with 33-1 or 62-1 at MOI 10 for 60 min. As controls BMNs were also infected with p32_08 (*exsA*^+^), p32_85 (*exsA*^T48I^) or an additional hyperactive *exsD*^T188P^ mutant (LU ExsD^T188P^) [33] (**Table 2**). IL-1β secretion was measured by ELISA and PMR was determined by LDH release assay. As compared to uninfected, IL-1β release was significantly increased upon infection with p32_85 (**Fig.7A**), likely due to ExoS-driven NLRP3 inflammasome activation [29]. IL-1β release in response to infection with the other isolates was variable but not significantly different from p32_85 (**Fig.7A**). PMR in BMNs infected with p32_85 was significantly increased compared to uninfected and p32_08 (**Fig.7B**) as previously shown [27]. PMR in BMNs infected with the hyperactive mutant 33-1 was similar to p32_85 while UL ExsD^T188P^ and 62-1 were lower (**Fig.7B**). CFU determination indicated that lower PMR by LU ExsD^T188P^ was not due to lower bacterial survival (**Fig.S5**).

**Fig.7.**
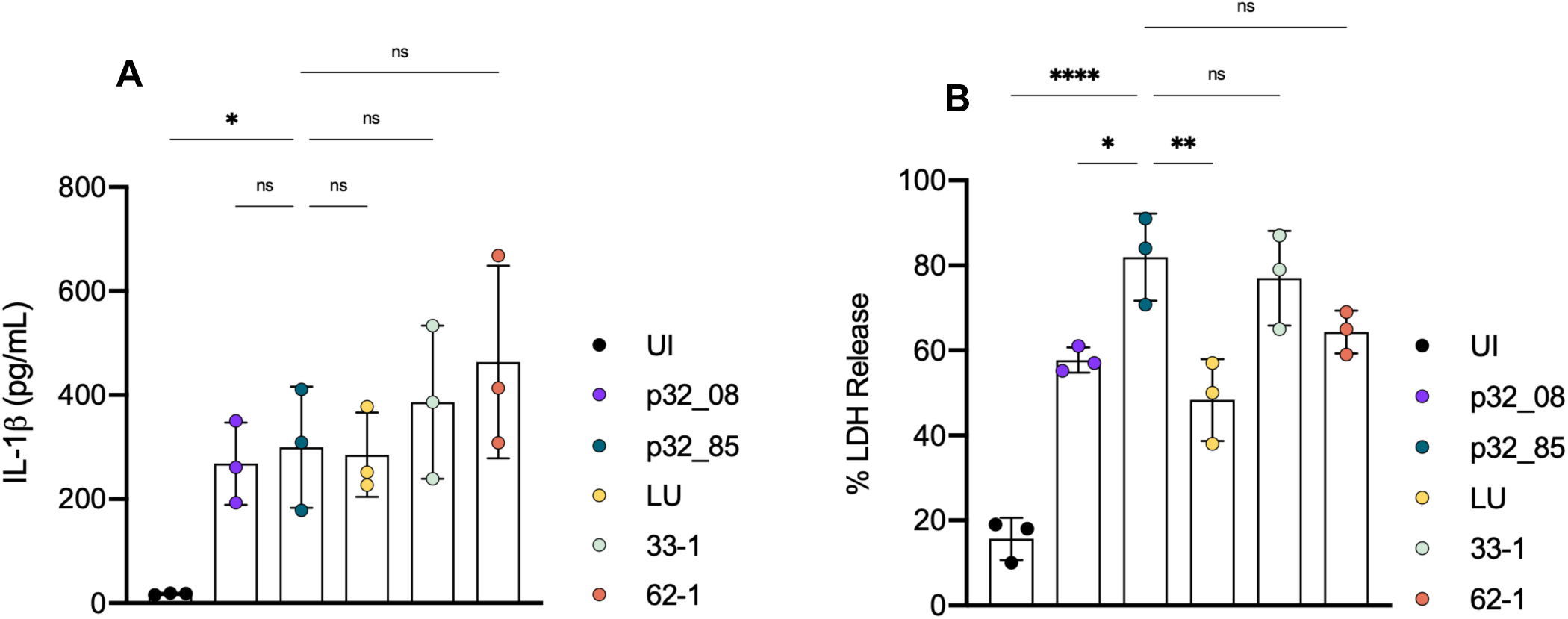
Analysis of IL-1β release and PMR in BMNs infected with *P. aeruginosa.* LPS-primed C57BL/6J BMNs were left uninfected (UI) or infected for 60 min with indicated strains at MOI 10 and analyzed for released IL-1β (A) or LDH (B). Data represent normalized values for 2.5x10^5^ cells/well ± the standard deviation from three independent experiments. Significant differences were determined by one-way ANOVA comparing to p32_85. ns, not significant; * P<0.05; ** P<0.01; **** P<0.0001. LU is strain LU ExsD^T188P^.

## Discussion

The use of bioinformatics to identify positively selected codons in genes from sequenced clinical isolates is a technically and conceptually innovative approach to identify and carry out structure/function analysis of bacterial virulence proteins [51, 52, 63]. As the number of sequenced clinical isolates increases, there is an important opportunity to leverage this type of analysis pipeline to understand the functions of products of virulence genes that undergo positive selection during infection. Results with *Salmonella* [51] were used to demonstrate that there is a context-dependent change in the fitness effect of the (in)organic phosphate antiporter *glpT* gene during *Salmonella* Typhimurium infection [53]. Our results with the *P. aeruginosa exs* genes provide an additional proof of principle for this concept.

From a stringent analysis of ∼24,000 *P. aeruginosa* genomes we identified 47 and 45 nonsense or missense codon changes in *exsA* and *exsD* respectively. These numbers far exceed the neutral expectations, indicating that most of these changes were positively selected. Furthermore, all four amino acid changes to ExsE affect the Arg-X-Val-X-Arg motifs involved in binding to ExsC, suggesting that they were selected because they disrupt this binding, producing a hyperactive T3SS phenotype. The four changes observed in *exsC* do not allow any conclusions about positive selection, though two of them—a change in the start codon and a premature stop codon—are expected to diminish or eliminate functional ExsC, diminishing T3SS function.

There are several possible explanations for why *exsA* and *exsD* are apparent “hotspots” for positively selected codon changes in this locus. In terms of gain of function mutations, codon changes in *exsA* and *exsD* may uniquely provide the maximal primed and loaded T3SS phenotype favored by positive selection. In addition, the impact of gain of function codon changes on the intrinsic growth rates of hyperactive mutants needs to be considered. Thus far we have not observed a measurable difference in the growth rates of p32_08 (*exsA*^+^) and p32_85 (*exsA*^T481^) in LB [27]. Lin et al. reported that a PA14 Δ*exsA* mutant grew slightly faster than a Δ*exsD* mutant in minimal media [45], indicating that, surprisingly, the hyperactive T3SS phenotype is not associated with substantial costs in bacterial replication rates in vitro.

Jorth et al. demonstrated that the LU *exsD*^T188P^ hyperactive T3SS mutant killed 90% of experimentally infected C57BL/6 mice within 24 hours as compared to 0% lethality in lung infections with the control *exsD*^+^ strain [33]. It was not determined is this hypervirulence was due to enhanced bacterial growth or exacerbated inflammation, or both. In addition, the virulence of the p32_85 *exsA*^T481^ hyperactive T3SS mutant had not been investigated in a mouse infection model. Our data show that p32_85 *exsA*^T481^ is hypervirulent in a CF mouse lung infection, causing significantly exacerbated weight loss as compared to the p32_85 *exsA*^+^ control. In addition, H&E staining of lung sections indicated a trend toward increased inflammatory cell recruitment to lungs infected with the hyperactive mutant. Organ burdens of the hyperactive mutant were not significantly elevated, suggesting that disease was caused by increased inflammation rather than increased bacterial growth. Hyperactive *P. aeruginosa* T3SS mutants can have a fitness advantage in the airways of pwCF, albeit in a location and or time dependent manner [27, 33-35]. Additional studies are needed to understand the variables that allow hyperactive *P. aeruginosa* T3SS mutants to have a fitness advantage in the airways of pwCF.

Excessive inflammatory responses to the T3SS are known to lead to worse outcomes to lung infection with *P. aeruginosa* [30, 32]. Excessive release of inflammatory molecules by macrophages and neutrophils in response to the hyperactive T3SS could contribute to effector-triggered pathogenesis [30, 32]. Increased macrophage detection of the hyperactive T3SS in the LU *exsD*^T188P^ strain results elevated IL-1β release and pyroptosis [33]. Enhanced ExoS-driven PMR in neutrophils as seen with p32_85 *exsA*^T481^ could result in danger-associated molecular pattern (DAMP) release [27]. The enhanced CitH3 and incomplete NET extrusion process detected in neutrophils infected with p32_85 *exsA*^T481^ appears to release certain DAMPs, e.g. HMGB1, but not histones [21, 27]. Other proinflammatory outcomes of incomplete NET extrusion remain to be defined. Measuring inflammatory responses of human CF neutrophils (lung and peripheral blood) after infection with *P. aeruginosa* hyperactive T3SS isolates is also important. Lung vs. peripheral neutrophils can have different phenotypes in CF [64] so a comparison of these two cell types is relevant.

*P. aeruginosa-*driven DAMP release in response to macrophage pyroptosis or neutrophil PMR is dependent on NINJ1 [21, 27]. Upon activation, monomeric NINJ1 localizes to the plasma membrane and oligomerizes to form ruptures large enough to release LDH [41-44]. Exposure of cells to 5 mM glycine prevents NINJ1 oligomerization and PMR [41]. Use of mice with conditional deletion of Ninj1 could be used to determine if DAMP release contributes to hypervirulence of p32_85 *exsA*^T481^. Alternatively, exogenous glycine supplementation has therapeutic potential to attenuate lung damage and virulence in mice challenged with hyperactive *P. aeruginosa* mutants, as suggested from previous experiments [65, 66].

Codon changes at position 48 of *exsA* do not appear to prevent interaction with, or negative regulation by, ExsD [27, 40] suggesting that ExsA^T48I^ is hyperactive for another reason. For example, an ExsA^T48A^ variant was shown to be negatively regulated by ExsD in an in vitro transcription assay [40]. In addition, ExsA^T48I^ showed no defect in ExsD interaction as shown by bacterial two-hybrid analysis [27]. However, the ExsA-ExsD binding mode is complex [37] and in vitro transcription or bacterial two-hybrid analysis may not always capture defective regulatory interactions between these proteins [40]. Position 48 is in the β-barrel, a domain which in AraC regulates a dimer conformation change to allow promoter occupancy [57]. How the β-barrel regulates ExsA is unknown, yet this domain appears to be a hotspot for positive selection of codon changes (18 total) in *exsA* during infection. AF3 modeling indicated that as compared to ExsA^+^ the ExsA^T48I^ dimer is in a different conformation that might facilitate receptor occupancy. Thus, ExsA^T48I^ may be constitutively active for this dimer conformation change. Confirming this model will require testing additional positively selected codon changes in *exsA* for hyperactivity, as well as measuring binding of the transcription factor to promoter DNA in vitro. These results will also provide new insights into regulation of ExsA by the β-barrel domain.

T3SS priming in *P. aeruginosa* is mediated by a free pool of ExsA produced under control of cAMP and Vfr [45, 49]. Our immunoblotting data show that ExsA^T48I^ does not increase production of itself, but does increase levels of ExsD, ExoS, and PcrV. In addition, ExsA^T48I^ appears to negatively regulate genes important for flagellar motility. Results of microscopy confirm that ExsA^T48I^ promotes a primed and loaded T3SS phenotype at the single cell level. Primed and loaded T3SSs could be critical for pathogenesis as they would increase the likelihood that *P. aeruginosa* has made an injectisome and effectors before encountering a neutrophil.

Based on the frequency of finding codon changes in *exsA* and *exsD* through searching ∼24,000 genomes it might appear that hyperactive mutants are rare (∼1:250). This prevalence figure may underestimate the pathogenic advantage of hyperactive T3SS *P. aeruginosa* strains for several reasons. First, the bioinformatic search for positively selected codon changes in *exs* genes in sequenced genomes in the NCBI Pathogens database was stringent and included quality control checks that may have missed some hyperactive mutants. Second, there may be few hyperactive mutant genomes in the database. Hyperactive mutants can exist transiently in pwCF, suggesting they have a pathogenic advantage in a time or location-dependent manner. Isolates from new infections, especially in young children with CF, may be enriched for hyperactive mutants, yet these genomes may be underrepresented in the database. In our analysis of 10 isolates from young children with CF five were T3SS positive (50%) and 2 were hyperactive (20%), with primed and loaded T3SS phenotypes similar to p32_85 *exsA*^T481^. The isolate 33-1 has a stop codon in *exsD* that truncates the protein, explaining the hyperactive phenotype. Although 10 is a small sampling size, these results raise the possibility that hyperactive T3SS *P. aeruginosa* strains are more prevalent in children with CF than previously known. Screening additional collections of isolates from children with CF (e.g. [67]) will test this possibility.

## Materials and Methods

### Analysis *P. aeruginosa* genomes in NCBI Pathogens Database for positively selected codon changes in *exs* genes

A recent build of the NCBI Pathogens data for P. aeruginosa, PDG000000036.1782, was analyzed. The sequence of the P. aeruginosa strain PAO1 chromosome, accession NC_002516.2, was used as a reference. Single-nucleotide changes that occurred within clusters of closely related isolates were inferred by maximum parsimony and mapped to the PAO1 ortholog as described in Cherry [51]. These changes were filtered by strict requirements for supporting sequence reads as described in Cherry [68]. The polarity of the *exs* amino acid replacements implied by the midpoint rooting of the phylogenetic trees was in all but one case consistent with the ancestral amino acid being that found in PAO1. For the single exception, in *exsB*, the rooting implied that the PAO1 amino acid was derived. As this could be explained by a slight misrooting of the tree, the PAO1 residue was assumed to be ancestral. The same assumption was used to polarize changes inferred to have occurred on branches descending directly from the root of the tree, which were included in the analysis. Neutral expectations were calculated as in Cherry [51, 52], except that the nearest neighbor model described [52] was not applied due to an insufficient number of nucleotide changes.

### Bacterial strains and growth conditions

*P. aeruginosa* strains used in this study are listed in Table 2. *P. aeruginosa* strains for neutrophil infections and secretion assays were grown on Luria-Bertani (LB) agar plates, and in LB high salt broth (11.7g/L NaCl) supplemented with MgCl_2_ (10mM) and CaCl_2_ (0.5mM) at 37°C [17]. *P. aeruginosa* for murine acute infection were grown in LB broth. The p*exoT*-sfGFP reporter was conjugated from *E. coli* into *P. aeruginosa* as described [45].

### Ethics statement

All experiments with mice were carried out in accordance with a protocol that adheres to the Guide for the Care and Use of Laboratory Animals of the National Institutes of Health (NIH) and was reviewed and approved (approval 2148) by the Institutional Animal Care and Use Committee at Dartmouth College. The Dartmouth College animal program is registered with the U.S. Department of Agriculture (USDA) through certificate number 12-R-0001, operates in accordance with Animal Welfare Assurance (NIH/PHS) under assurance number D16-00166 (A3259-01) and is accredited with the Association for Assessment and Accreditation of Laboratory Animal Care International (AAALAC, accreditation number 398).

### Mouse strains

The F508del mice (also referred to as *Cftr*^F508del^ or *Cftr*^em1Cwr^) were obtained from Craig Hodges at the Case Western Reserve University CF Mouse Model Core [55]. These mice have a precise deletion inactivating *Cftr* codon F508 on the C57BL/6 background. Although F508del mice have a milder CF phenotype as compared to *Cftr* knockouts, they can be maintained without laxatives that might disrupt the microbiota and breed well in the homozygous mutant background. C57BL/6J mice from Jackson Laboratories used for BMNs.

### Infection of F508del mice with p32_08 or p32_85

For acute infections with p32_08 and p32_85, overnight (16-hour) cultures of *P. aeruginosa* were sub-cultured 1:100 in fresh LB and grown at 37°C to mid-log phase at OD_600_= 0.5. *Cftr^F508del^* mice were anesthetized by isoflurane inhalation and oropharyngeally challenged with 5x10^5^ CFUs in 100μl PBS. Mice were monitored for weight loss and survival for 5 days and then sacrificed for CFU analysis. Mice were euthanized by CO_2_ inhalation. Lung homogenate was collected following full lung dissection, separation into left and right, fixation of right lungs and manual homogenization of left lungs using 2mL of cold PBS. Lung homogenate was used for CFU assay by plating serial dilutions on LB agar or *Pseudomonas* Isolation Agar plates, incubation at 37C. Right lungs were processed for H&E staining.

### Analysis of T3SS protein production and secretion by immunoblotting

Overnight (16-hour) cultures of *P. aeruginosa* were sub-cultured 1:100 in fresh LB high-salt broth supplemented with MgCl_2_ (10mM) and CaCl_2_ (0.5mM) with or without EGTA (5mM) and grown at 37°C to mid-log phase at OD_600_= 0.5. 1ml of culture was pelleted, supernatant was mixed with trichloro acetic acid (final concentration 10%) and incubated overnight at 4°C with shaking. Proteins were pelleted by centrifugation, washed with acetone and resuspended in 1x sample buffer containing DTT. Bacterial pellets were lysed in Bugbuster and soluble lysates were obtained by centrifugation. Secreted proteins and cell lysates were run on 4-12% NuPAGE Bis-Tris SDS-PAGE gels (Invitrogen by ThermoFisher Scientific) and transferred to PVDF membranes (ThermoFisher Scientific) using an iBlot 2 Gel Transfer Device (Life Technologies). Membranes were blocked in 5% non-fat dairy milk and incubated with primary antibody overnight. The primary antibodies used were rabbit polyclonal specific for ExoS or PcrV (from Arne Rietsch) or ExsA or ExsD (from Tim Yahr and Matt Wolfgang). HRP-conjugated anti-rabbit (Jackson Immuno Research) was used as a secondary antibody. Protein bands reacting with antibodies were visualized using chemiluminescent detection reagent (GE Healthcare) on an iBright FL1500 (ThermoFisher Scientific).

### Analysis of T3SS bistability by microscopy

Overnight (16-hour) cultures of *P. aeruginosa* expressing p*exoT*-sfGFP [45] were sub-cultured 1:100 in fresh LB high-salt broth supplemented with MgCl_2_ (10mM) and CaCl_2_ (0.5mM) with or without EGTA (5mM) and grown at 37°C to mid-log phase at OD_600_= 0.5. 1ml of culture was pelleted and resuspended in 4% PFA for 30min. Pellets were washed with PBS and resuspended in Hoechst 33342 (2μg/ml) and incubated for 30min. Pellets were washed with PBS and resuspended in 100μl of PBS and plated on agarose pads mounted on a microscope slide and then covered over with a cover slip. Images were obtained using a Nikon C2 confocal system and an Apo 40x/1.25 NA water objective. Images were processed using FIJI software with custom-made macros. These macros as well as other relevant data underlying this research is available in a public data repository (Zenodo 10.5281/zenodo.23603675).

### Flagellar motility assays

Swim assays were performed according to previously published methods [69]. Semisolid swim assay media was formulated with an agar concentration of 0.3%. To control experimental variability, parameters for nutrient medium volume, plate thickness, and plate dryness were standardized during pouring and curing. Overnight (16-hour) cultures of *P. aeruginosa* p32_08 and p32_85 were sub-cultured 1:100 in fresh LB high-salt broth supplemented with MgCl_2_ (10mM) and CaCl_2_ (0.5mM) with or without EGTA (5mM) and grown at 37°C to mid-log phase at OD_600_= 0.5. Plates were centrally inoculated on or just below the surface with a fixed volume of the fresh bacterial culture to initiate a localized colony zone before being immediately transferred to 37°C and incubated faceup. Representative images of plates were captured with a camera. Total surface coverage area and branching morphology were visually assessed. Surface area was quantified on plates incubated faceup using ImageJ by manually outlining and measuring the whole colony for surface area. Colony diameter was quantified by measuring the shortest line through the point of inoculation from one edge of the colony to the opposite edge.

### AlphaFold3 modeling

Structural modeling of the ExsA, ExsA^T48I^, and ExsA^N18S^ dimers was performed using the AlphaFold3 webserver [58]. Highest confidence models were visualized using ChimeraX (https://pubmed.ncbi.nlm.nih.gov/28710774/). Structure overlap between ExsA and ExsA^N18S^ was performed using the included MatchMaker function with the ExsA dimer model as the reference structure.

### Preparation of Neutrophils

Bone marrow was isolated from tibia and femur exudates of 8-12 week old C57BL/6J mice. Single-cell suspensions of bone marrow were prepared, and neutrophils were isolated following Miltenyi MACS Neutrophil Isolation Kit. Neutrophils were plated at a density of 2.5x10^5^ in 100μL of OptiMEM in a 96 well plate. The neutrophils were primed with 100ng/mL O26:B6 *Escherichia coli* LPS (Sigma) and incubated overnight (16h) at 37°C with 5% CO_2_.

### Neutrophil Infection Assays

Overnight (16-hour) cultures of *P. aeruginosa* were sub-cultured 1:100 in fresh LB high-salt broth supplemented with MgCl_2_ (10mM) and CaCl_2_ (0.5mM) and EGTA (5mM) and grown at 37°C to mid-log phase at OD_600_= 0.5. Cultures were then pelleted, the LB removed, and bacteria were resuspended in PBS to the original volume. Bacterial suspensions were then diluted to an MOI of 10 or 1 in OptiMEM. Overnight media from the neutrophils was removed and replaced with 100μL of OptiMEM containing bacteria. Plates were then incubated at 37°C with 5% CO_2_ for 60, 120, or 180 minutes. Cell supernatants (MOI 10) were collected for cytokine ELISAs and lactate dehydrogenase (LDH) assays. Cell supernatants were removed (MOI 1), and cells were lysed with 100μL 0.1% NP-40 and serially diluted in PBS and plated on LB agar for CFU assays.

#### IL-1β quantification

Murine IL-1β in neutrophils supernatants was quantified using ELISA kits (R&D Systems, MLB00C) following the manufacturer’s instructions.

#### LDH quantification

LDH in neutrophil supernatants was quantified using the CytoTox 96 Non-Radioactive Cytotoxicity Assay (Promega) following the manufacturer’s instructions.

#### Statistical analysis of data

Experimental data generated from at least three independent experiments are analyzed for significance using GraphPad Prism software. Probability (*p*) values for ELISA and LDH data are calculated using one-way analysis of variance (ANOVA) or grouped two-way ANOVA with Tukey’s posttest. P values of less than or equal to 0.05 were considered significant. Mouse weights were determined for significance by multiple T-test and a multiple-comparisons correction (Holm–Šídák). Right lung lobe CFU were evaluated for significance by Mann Whitney test. GFP MFI data groups were compared by a mixed model paired t-test with Bonferroni-Holm correction for multiple comparisons.

## Acknowledgements

We thank members of the Bliska Laboratory for providing feedback on this manuscript. We also thank Wei Wang for assistance with mouse husbandry, Aurora Golden for the H&E images of lungs from uninfected F508del mice, Arne Rietsch for the PcrV and ExoS antibodies and the PAO1F and PAO1F Δ*pscD* strains, Tim Yahr and Matt Wolfgang for the ExsA and ExsD antibodies, Dan Aridgenes and Deborah Hogan for the DH1137 strain, Luke Hoffman for the clinical isolates from children with CF and Barbara Kazmierczak for the p*exoT*-GFP reporter, bacterial strains and for advice on bistability assays.

Supported by a grant from the National Institutes of Health (the Dartmouth Cystic Fibrosis Training Program (T32HL134598 to ADR)), the Cystic Fibrosis Foundation Research Development Program (BOMBER24G0 to THH), and funds from the Philip Hanlon and Gail Gentes Cluster for Personalized Treatments for Cystic Fibrosis.

This research was also supported in part by the Intramural Research Program of the National Institutes of Health (NIH). The contributions of the NIH author are considered Works of the United States Government. The findings and conclusions presented in this paper are those of the author and do not necessarily reflect the views of the NIH or the U.S. Department of Health and Human Services.

---

**Fig.S1.**
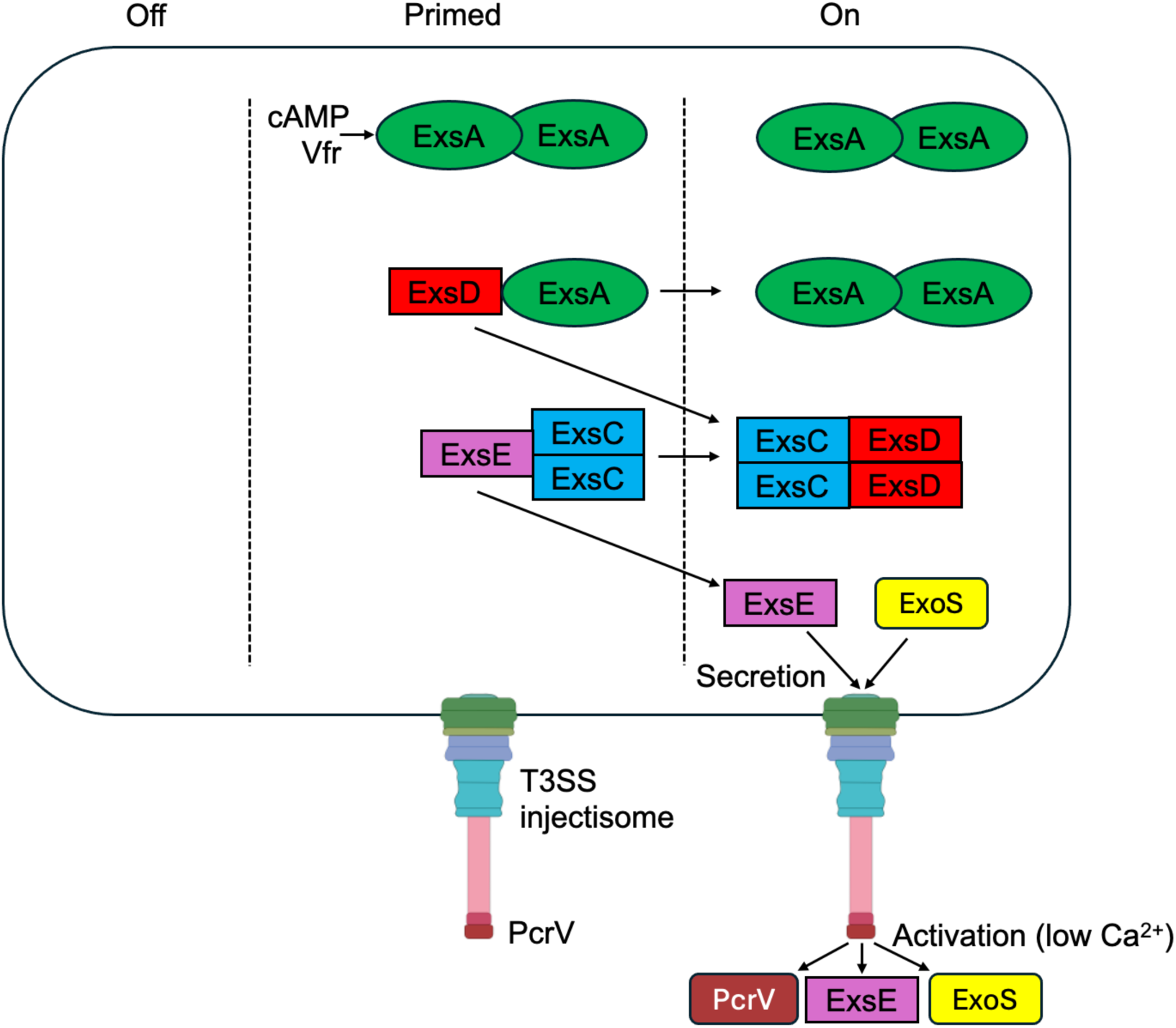
Model of *P. aeruginosa* bistability and partner switching mechanism of T3SS regulation. Products of the *exsCEBA* operon and *exsD* contribute to regulation of bistability and connect regulation of T3SS gene expression to secretion. In the absence of secretion activating signals such as low Ca^2+^ or host cell contact *P. aeruginosa* cells in a population are in the off or primed condition. In the off state ExsA production is below detectable levels. In the primed state sufficient free ExsA produced under control of cAMP-activated Vfr is available to promote expression of the ExsD, ExsE and ExsC regulators and the other operons leading to assembly of a functional T3SS injectisome with PcrV at the tip. In the on state in response to activating signals ExsE is secreted through the injectisome, releasing ExsC to sequester ExsD, which frees ExsA to activate maximal T3SS expression, secretion of PcrV and production and secretion of ExoS.

**Fig.S2.**
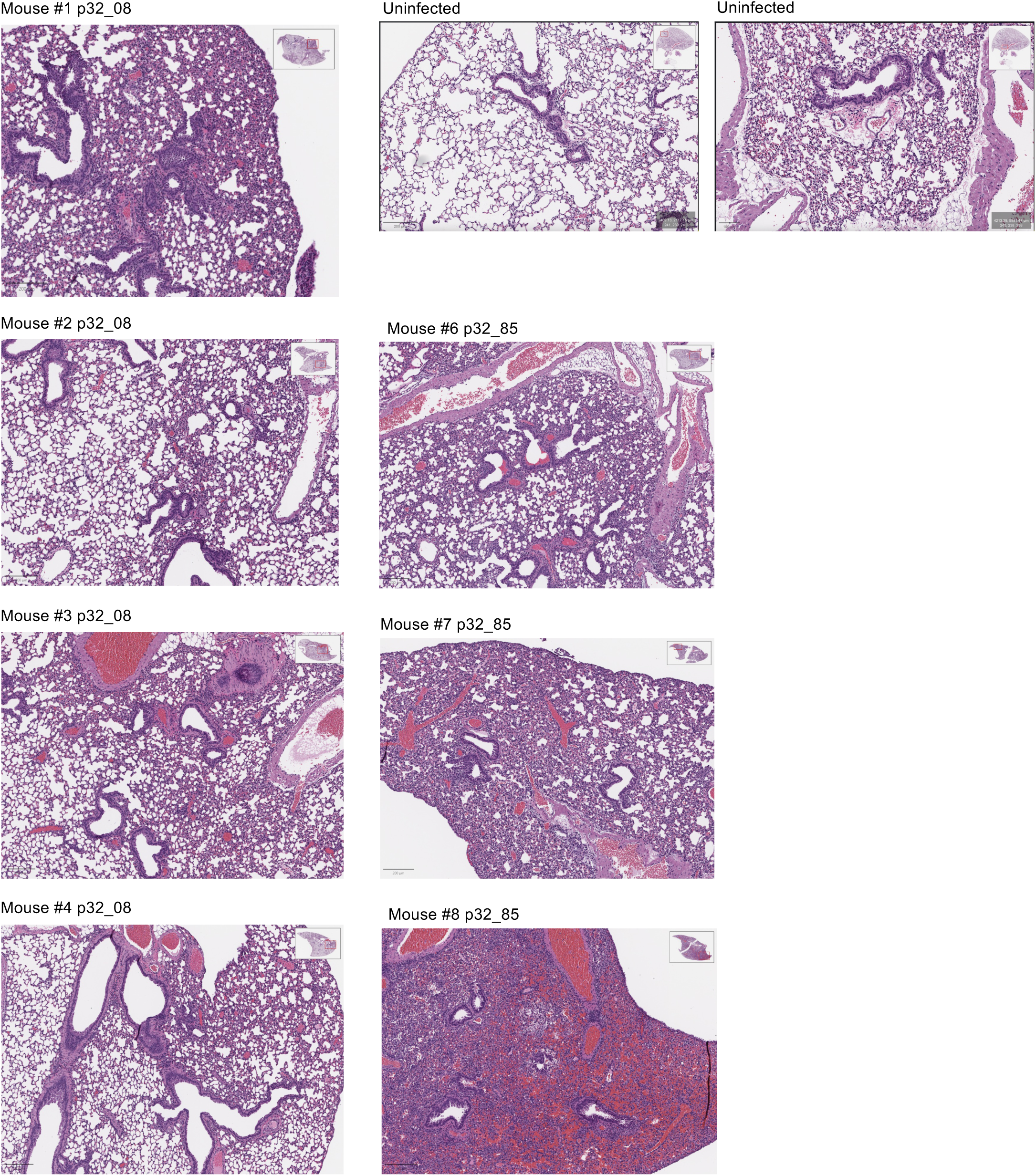
Lung H&E from F508del mice left uninfected or infected with p32 isolates 08 (*exsA*^+^) or 85 (*exsA*^T48I^). Representative H&E images of left lobe lung sections at 120 h and 10x magnification from the third independent infection experiment. Representative images of left lobe lung sections from an uninfected mouse at 10x magnification are shown in upper right. Insets in upper right corners show full lung images. Scale bars are 200 μM.

**Fig.S3.**
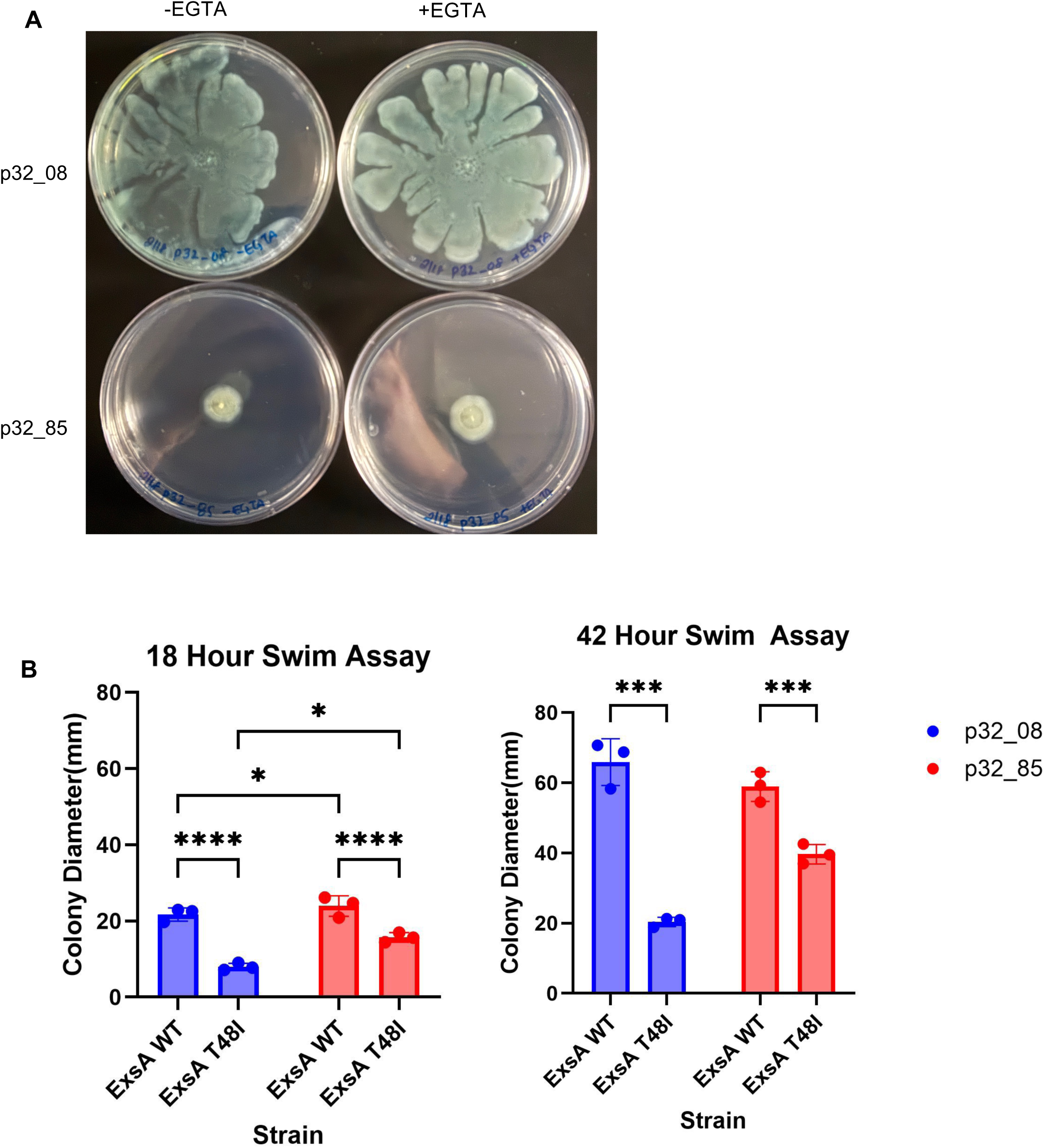
Swim assay comparison of p32_08 and p32_85. (A) p32_08 or p32_85 were grown in LB in the absence or presence of EGTA and spotted onto swim agar plates and incubated at 37°C faceup. Representative images after overnight (18hr) incubation are shown. (B). The indicated strains were grown in the absence of EGTA and assayed for swim motility faceup after 18 or 42 hr. Colony diameter was quantified using ImageJ. Results are pooled from 3 independent experiments and analyzed for significance using one-way ANOVA * P<0.05; *** P<0.001; **** P<0.0001.

**Fig.S4.**
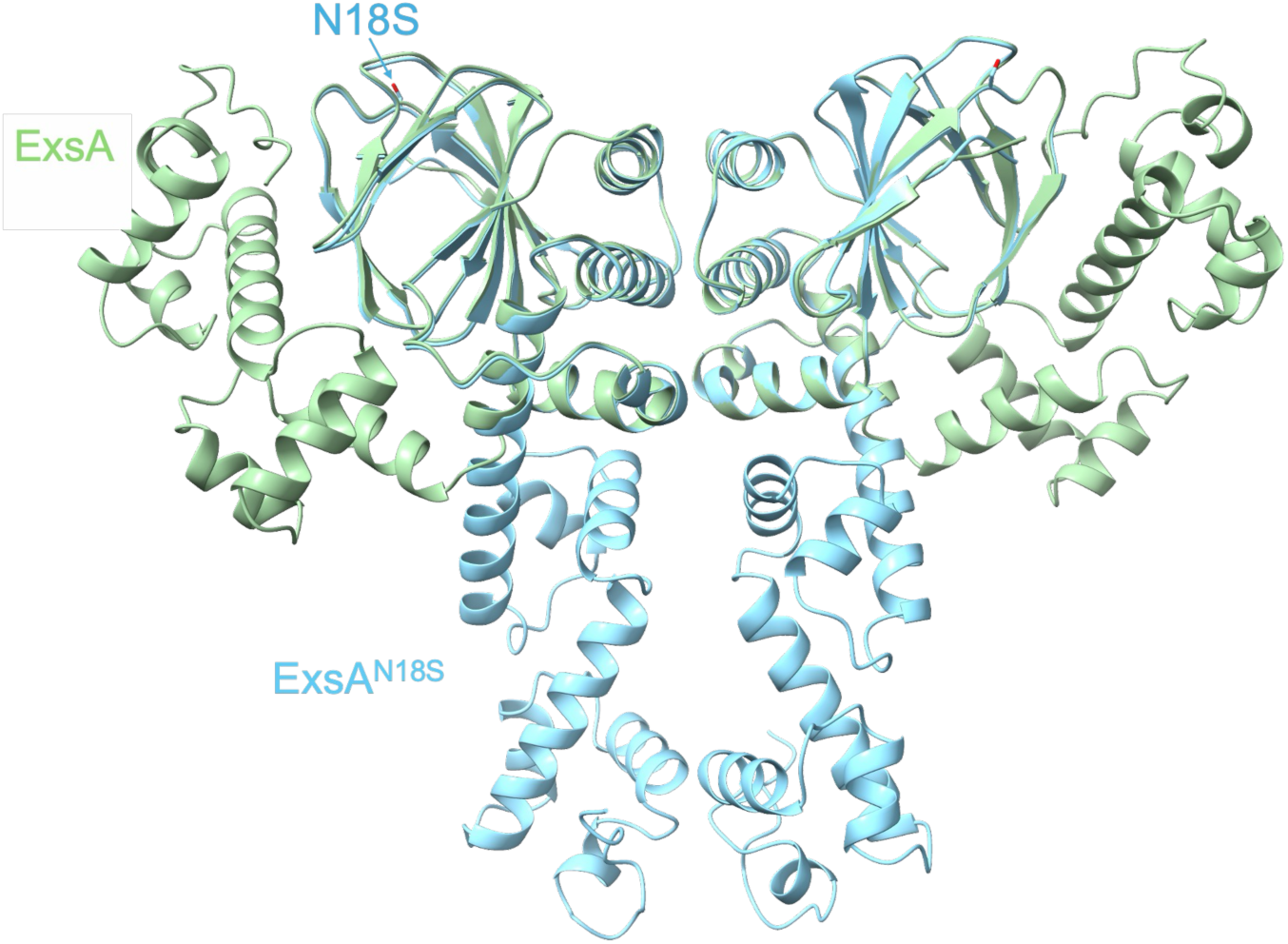
AF3 models of ExsA and ExsA^N18S^. ExsA (green) and ExsA^N18S^ (blue) dimers superimposed to show conformational change in HTH domains resulting from N18S change. In AF3 the Predicted Template Modeling Score (pTM) measures the accuracy of the overall predicted structure of an individual chain or a whole complex. Scores above 0.5 indicate a likely correct overall fold. The pTM values for dimer structures of ExsA and ExsAN18S are 0.66 and 0.57 respectively.

**Fig.S5.**
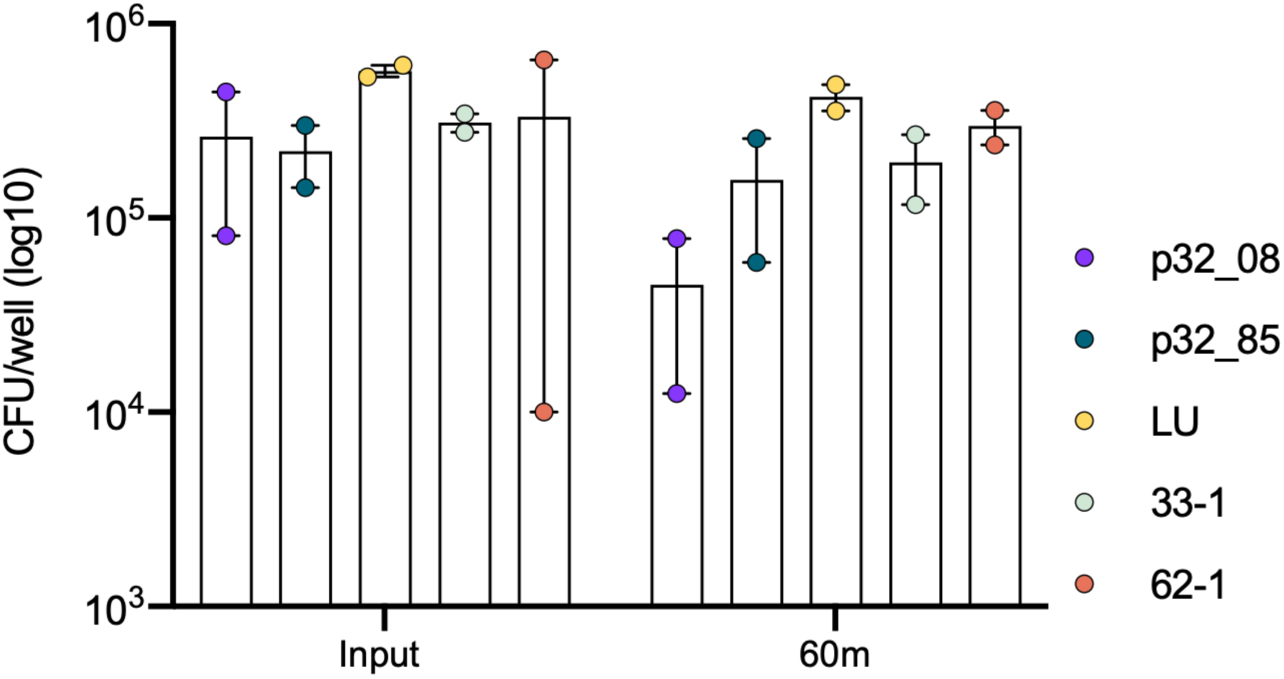
CFU analysis in BMNs infected with *P. aeruginosa.* BMNs were infected with the indicated strains at an MOI of 1 for 60 min. Samples of bacteria inoculated into each well (Input, T=0 min) and total bacterial-neutrophil co-cultures solubilized in 0.1% NP-40 detergent (T=60 min) were processed by serial dilution and plating to determine CFUs. Data represent normalized values for 2.5x10^5^ cells/well ± the standard deviation from two independent experiments

## Notes

### Competing Interest Statement

The authors have declared no competing interest.

### Summary of Updates

This version of the manuscript has been revised to update supplemental figure labels.

